# Enhancer-promoter contact formation requires RNAPII and antagonizes loop extrusion

**DOI:** 10.1101/2022.07.04.498738

**Authors:** Shu Zhang, Nadine Übelmesser, Mariano Barbieri, Argyris Papantonis

**Affiliations:** Institute of Pathology, University Medical Center Göttingen, 37075 Göttingen, Germany

**Keywords:** chromosome conformation capture, RNA polymerase, loop extrusion, Polycomb complex

## Abstract

Mammalian chromosomes are folded by converging and opposing forces. Here, we tested the role of RNAPII across different scales of interphase chromatin folding in a cellular system allowing for its auxin-mediated degradation. We used Micro-C and computational modeling to characterize subsets of loops differentially gained or lost upon RNAPII depletion. Gained loops, extrusion of which was antagonized by RNAPII, almost invariably formed by engaging new or rewired CTCF anchors. Lost loops selectively concerned contacts between enhancers and promoters anchored by RNAPII. Surprisingly, promoter-promoter contacts were almost insensitive to polymerase depletion and sustained cohesin occupancy in its absence. Selective loss of enhancer-promoter contacts explains the repression of most genes. Together, our findings reconcile the role of RNAPII in transcription with that in setting-up regulatory 3D chromatin architectures genome-wide, while also revealing a direct impact on cohesin loop extrusion.

## INTRODUCTION

Genomic functions, such as gene expression and DNA replication, require a tunable three-dimensional (3D) architecture of interphase chromatin (Aboelnour & Bonev, 2021; Razin & Kantidze, 2022). Work over the last decade, combining genome-wide chromosome conformation capture assays with the removal of different chromatin components, has attributed most hallmarks of this 3D architecture to the interplay between two factors, the insulator protein CTCF (Xiang & Corces, 2021) and the ring-shaped cohesin complex (van Ruiten & Rowland, 2021). 3D chromatin domains [from the Mbp-sized “topologically-associating domains” (TADs; Beagan & Phillips-Cremins, 2020) to kbp-sized “loop domains (Rao et al., 2014)] are insulated from one another via CTCF-demarcated boundaries, while the chromatin at and within these domains is actively extruded into loops by cohesin (Fudenberg et al., 2016; Davidson et al., 2019; Kim et al., 2019). Removing CTCF from chromatin leads to the loss of insulation at domain boundaries (Nora et al., 2017), while cohesin removal leads to the elimination of CTCF-anchored loops (Rao et al., 2017; Wutz et al, 2017). Physical interaction of an extruding cohesin-complex ring with two convergently-oriented, yet distal, CTCF-bound sites determines loop length and stabilizes its formation together with STAG cofactors (Rao et al., 2014; Li et al., 2020; Wutz et al., 2020).

Loops anchored at CTCF-bound sites appear as prominent dot-like features off the diagonal of high-resolution Hi-C contact maps (Rao et al., 2014). These dots disappear in cells where the cohesin-loading factor, NIPBL, is eliminated (Schwarzer et al., 2017), but multiply in cells lacking the cohesin-release factor WAPL (Haarhuis et al., 2017). Thus, loop formation requires efficient cohesin loading onto chromatin, while its processivity is restricted by regulated “load-unload” cycles. However, recent live-cell imaging of the mouse *Fbn2* locus showed that full looping is rarely achieved and that, most of the time, cohesin-extruded loops within an active domain form without bringing both CTCF boundaries together (Gabriele et al., 2022). This can be explained by the notion that 3D genome architecture results from the antagonistic interplay between loop extrusion and compartmentalization of homotypic (i.e., active-to-active or inactive-to-inactive) chromatin domains (Nuebler et al., 2018; Rada-Iglesias et al., 2018). For transcriptionally-active domains especially, patterns of activity constitute a good predictor of spatial chromatin partitioning as mapped by Hi-C (Ulianov et al., 2016; Rowley et al., 2017). Thus, one would predict that RNA polymerase II (RNAPII), a potent molecular motor capable of translocating and bridging DNA (Papantonis & Cook, 2011; Lee & Blobel, 2016), influences 3D genome architecture via both physical interactions (Cook & Marenduzo, 2018) and transcription (Racko et al., 2018).

Studies in this direction have shown that chemical inhibition of transcription weakens insulation at TAD boundaries (Hug et al., 2017; Barutcu et al., 2019), while RNase A treatment of cell nuclei does not affect TAD structure, but does eliminate specific enhancer-promoter contacts (Brant et al., 2016; Barutcu et al., 2019). On the other hand, Hi-C maps generated upon depletion of Mediator complex subunits or upon inhibition of RNAPII elongation while re-expressing cohesin in RAD21-depleted cells had no discernible effect on CTCF loop formation (Rao et al., 2017; El Khattabi et al., 2019). This held true also when depleting the basal transcription factor TAF12 or RNAPII and using promoter-capture Hi-C or Hi-C, respectively (Jiang et al., 2020; Sun et al., 2021). Moreover, haploid human cells depleted of Mediator could not sustain a transcription-permissive chromatin architecture (Haarhuis et al., 2022). However, even kbp-resolution Hi-C contact maps do not prominently feature loops other than those anchored by CTCF. This was remedied by the introduction of Micro-C, a Hi-C variant using micrococcal nuclease (MNase) to fragment chromatin and reveal tens of thousands of transcription-based loops in mammalian chromosomes at sub-kbp resolution (Hsieh et al., 2020; Krietenstein et al., 2020). In fact, a capture-based adaptation of this approach, Micro-Capture-C (MCC), allowed mapping of 3D contacts between different *cis*-regulatory elements at near-base-pair resolution (Hua et al., 2021).

This and other data highlight the need for using Micro-C or MCC to conclusively dissect whether and how core components of the transcriptional apparatus, like RNAPII and Mediator (Cramer, 2019), contribute to the formation of 3D chromatin contacts. We did this by applying Micro-C on a diploid human cell line that allows for the auxin-mediated degradation of the largest RNAPII subunit, RPB1 (Nagashima et al., 2019). This way, we identified transcription-anchored loops, which we then grouped according to their response to RNAPII depletion. We combined experiments with *in silico* modeling to interpret the interplay between cohesin loop extrusion and RNAPII-mediated looping. In parallel to our study, an orthogonal one applying MCC to mouse cells acutely depleted of Mediator generated similar observations across different genomic loci (Preprint: Ramasamy et al., 2022).

## RESULTS

### Transcription-level architecture of human chromosomes

Previously, we used *in situ* Hi-C to identify changes in 3D genome architecture upon depletion of RNAPII from human diploid cells (Zhang et al., 2021). For these Hi-C experiments, we used non-synchronized G1-sorted DLD-1 cells allowing for the quantitative degradation of RNAPII upon auxin addition for 14 h (Nagashima et al., 2019). Although effects at the level of TADs and compartments were small (as also observed upon RNAPII removal from asynchronous mESCs; Jiang et al., 2020), we did identify ∼800 CTCF loops that emerged in RNAPII-depleted cells and were significantly larger than the loops in untreated cells (Zhang et al., 2021). However, interactions between RNAPII-bound sites were scarce in this data, hence our incomplete understanding of how RNAPs contribute to 3D chromatin folding.

To address this and obtain a comprehensive view of the transcription-based architecture of human cells, we performed Micro-C. We generated contact maps with just <1 billion pairwise contacts, which revealed fine intra-domain architecture compared to previous Hi-C data (despite similar sequencing depth; **Figure 1A,B**). This detailed view of 3D chromatin folding allowed detection of ∼30,000 loops, encompassing >80% of the loops detected by Hi-C (**Figure 1C**). The anchors of these loops mapped predominantly to the A (active) compartment, suggesting that multiple RNAPII-anchored loops were detected (**Figure 1D**). Indeed, when we stratified these ∼30,000 loops according to the presence or absence of transcription marks/RNAPII or CTCF at each of their anchors, we found that ∼25% could be classified as transcription-anchored, which increases to 40% once we add anchors that have CTCF in addition to RNAPII/H3K27ac. Moreover, Micro-C contact maps also feature the ∼8,200 loops that are shared with Hi-C more prominently (**Figure 1E**). Finally, our highly resolved 3D contact data allowed detection of thousands of instances where directional loop extrusion gave rise to stripes emanating by one CTCF and one transcriptional anchor (**Figure 1F**). Thus, Micro-C now allows us to ask how such transcription-anchored 3D interactions are remodeled upon RNAPII depletion.

**Figure 1.**
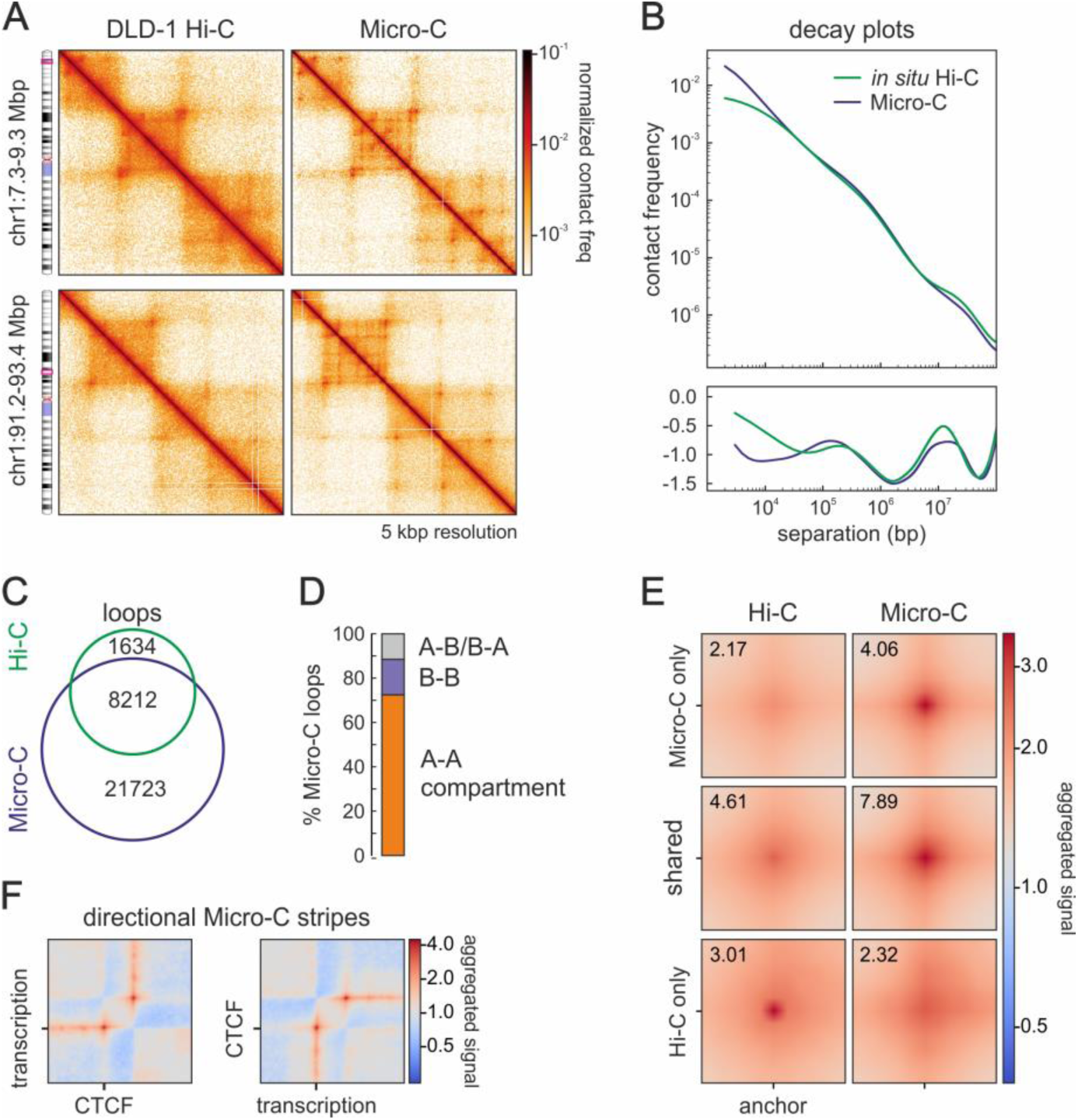
Micro-C enhances high-resolution views of 3D genome folding. (**A**) Comparison of Hi-C (left) and Micro-C 5-kbp resolution contact maps (right) from DLD1-mAID-RPB1 cells. (**B**) Hi-C (green) and Micro-C (blue) interaction frequency decay plots as a function of genomic distance (top) and their first derivative (bottom). (**C**) Venn diagram showing overlap of loops detected in Hi-C (green) and Micro-C contact maps (blue). (**D**) Bar plot showing per cent of Micro-C loops with anchors in the A- or B-compartment. (**E**) Aggregate plots showing Hi-C (left) and Micro-C signal (right) around shared and unique loops. (**F**) Average plots showing mean Micro-C stripe signal at loops with one CTCF and one transcriptional anchor.

### New CTCF-anchored loops emerge upon RNAPII depletion

To address how RNAPII affects 3D interactions, we used the 14-h auxin treatment that was determined previously (Zhang et al., 2021) and generated Micro-C data in the presence or absence of RNAPs. Under these conditions, the vast majority of RNAPII is removed from chromatin leading to decreased H3K27ac and cohesin levels in CUT&Tag experiments (**Figure S1A,B**). Following RNAPII depletion, we used Micro-C data to assess changes in nucleosome positioning. For example, nucleosomes around CTCF sites become markedly more ordered; this does not come at the expense of CTCF binding to cognate motifs, but does result in more focal ATAC-seq signal, i.e. to constrained accessibility (**Figure S1C**).

We next surveyed Micro-C contact profiles to discover a widespread emergence of new and longer loops. These new loops typically arise in and around domains that contain active genes which are, thus, silenced upon RNAPII depletion (**Figures 2A** and **S1D**). The >8,700 loops gained in the absence of RNAPII are significantly longer than both CTCF- and transcription-anchored loops in control cells (**Figure 2B**). They produce new positions of increased insulation and involve at least one CTCF-bound anchor, which is different from what we documented for loops lost upon RNAPII depletion (**Figure 2C-E**).

**Figure 2.**
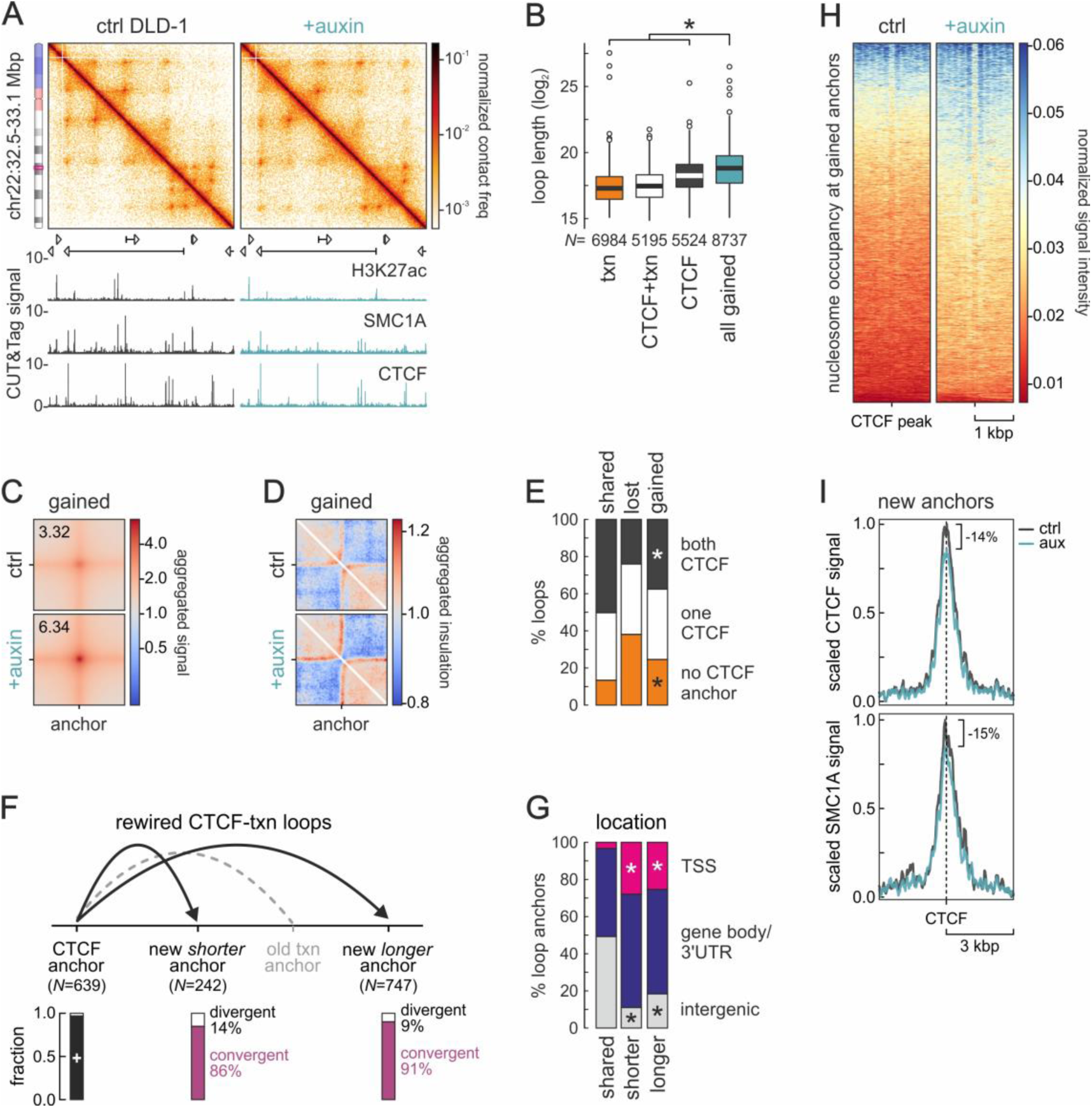
Loops forming upon RNAPII degradation engage new CTCF anchors. (**A**) Micro-C 2-kbp resolution contact maps from control (left) and auxin-treated cells (right) showing gain of loops in an exemplary genomic region on chr22 aligned to H3K27ac, CTCF, and SMC1A CUT&Tag tracks. (**B**) Box plots of the lengths of control loops with transcription-(orange), CTCF-(black) or transcription/CTCF-anchors (white) and loops gained upon auxin treatment (green). *: *P*<0.01, Wilcoxon-Mann-Whitney test. (**C**) Aggregate plots for loops gained in auxin-treated cells. (**D**) Plots showing mean insulation at the anchors of loops gained in auxin-treated cells. (**E**) Bar plots showing per cent of loops with no (orange), one (white) or two CTCF anchors (black) that are lost or gained in auxin-treated cells. *: different to shared loops; *P*<0.01, Fischer’s exact test. (**F**) Bar plots showing the fraction of convergent versus divergent CTCF motifs in gained loop anchors (*N*). (**G**) Bar plots showing per cent of gained loop anchors located in different genomic locations. (**H**) Heatmaps showing nucleosome occupancy around gained CTCF anchors from panel F. (**I**) Line plots showing mean SMC1A signal in the 6 kbp around gained loop anchors from panel F.

Looking into loops that have one anchor classified as “CTCF only” (e.g., the left one) and the other as “RNAPII only” (e.g., the right one), we found that they specifically rewire the latter. From a total of 989 such loops, 76% rewire to a new anchor further up/downstream that almost invariably contains a CTCF-bound site (**Figure 2F**). This often gives rise to nested loop structures (i.e., 639 unchanged loop anchors give rise to 989 new loops; **Figure 2A**). The orientation of CTCF motifs in these new anchors is predominantly convergent in respect to that in the unchanged anchor (**Figure 2F**). Interestingly, these new CTCF anchors are disproportionately located at TSSs and gene bodies, which become depleted of active RNAPs (**Figure 2G**). In the absence of RNAPII binding and transcription, nucleosomes around these anchors become more canonically spaced (**Figure 2H**), but show negligibly lower SMC1A levels (**Figure 2I**), despite the genome-wide drop in cohesin occupancy in RNAPII-depleted cells (see SMC1A signal in **Figures 2A** and **S1B-C**).

Finally, we asked what fraction of the loops gained in the absence of RNAPII form via H3K27me3-mediated interactions (there were hints of ∼200 such loops in our previous Hi-C data; Zhang et al., 2021). Although we saw no discernible changes in H3K27me3 CUT&Tag profiles from control and auxin-treated cells, ∼1,000 new loops arise that have H3K27me3 peaks in at least one anchor. This increases to 1,405 if we consider anchors with H3K27me3 peaks in the next (5- or 10-kbp) bin (**Figure S1F,G**). Such new loops typically emerge in bundles within facultative heterochromatin domains, often without CTCF contribution (**Figure S1E**). Therefore, our Micro-C data now explain the emergence of thousands of new and longer loops after RNAPII depletion from chromatin.

### RNAPII depletion leads to the selective loss of enhancer-anchored loops

Next, we asked which loops and contacts are lost upon RNAPII depletion. Lost loops were almost always found within CTCF loop-domain or TAD structures (**Figures 3A** and **S2A**). This is in line with transcription-anchored loops being the smallest (**Figure 2B**), with the overall reduced contact frequency between loci separated by <1 Mbp (**Figure 3B**), and with the notion that gene regulatory domains are encompassed by CTCF loops. Moreover, the anchors of lost loops were significantly less likely to contain CTCF than those of unchanged loops, and <25% of them have CTCF bound to both anchors (**Figure 3C**). Following stratification of loops into promoter-promoter (P-P) or enhancer-enhancer/promoter ones (E-E/E-P), we discovered that enhancer-anchored ones were selectively lost upon RNAPII depletion (**Figure 3D**), irrespective of whether they contained CTCF in either anchor (**Figure S2B**). This selective loss was also reflected in the strongly reduced cohesin occupancy of E-E/E-P compared to P-P loops (that was even more pronounced at the 590 super-enhancers; **Figures 3E** and **S2C**). Surprisingly, much like their looping propensity, H3K27ac levels at promoters remained unaffected, while those at enhancers dropped by >50% (**Figure S2D**). In total, >2,200 E-P loop domains dissolve upon RNAPII depletion and they harbor almost half (650/1360) of all genes significantly downregulated following auxin treatment (RNA-seq data from Zhang et al., 2021).

**Figure 3.**
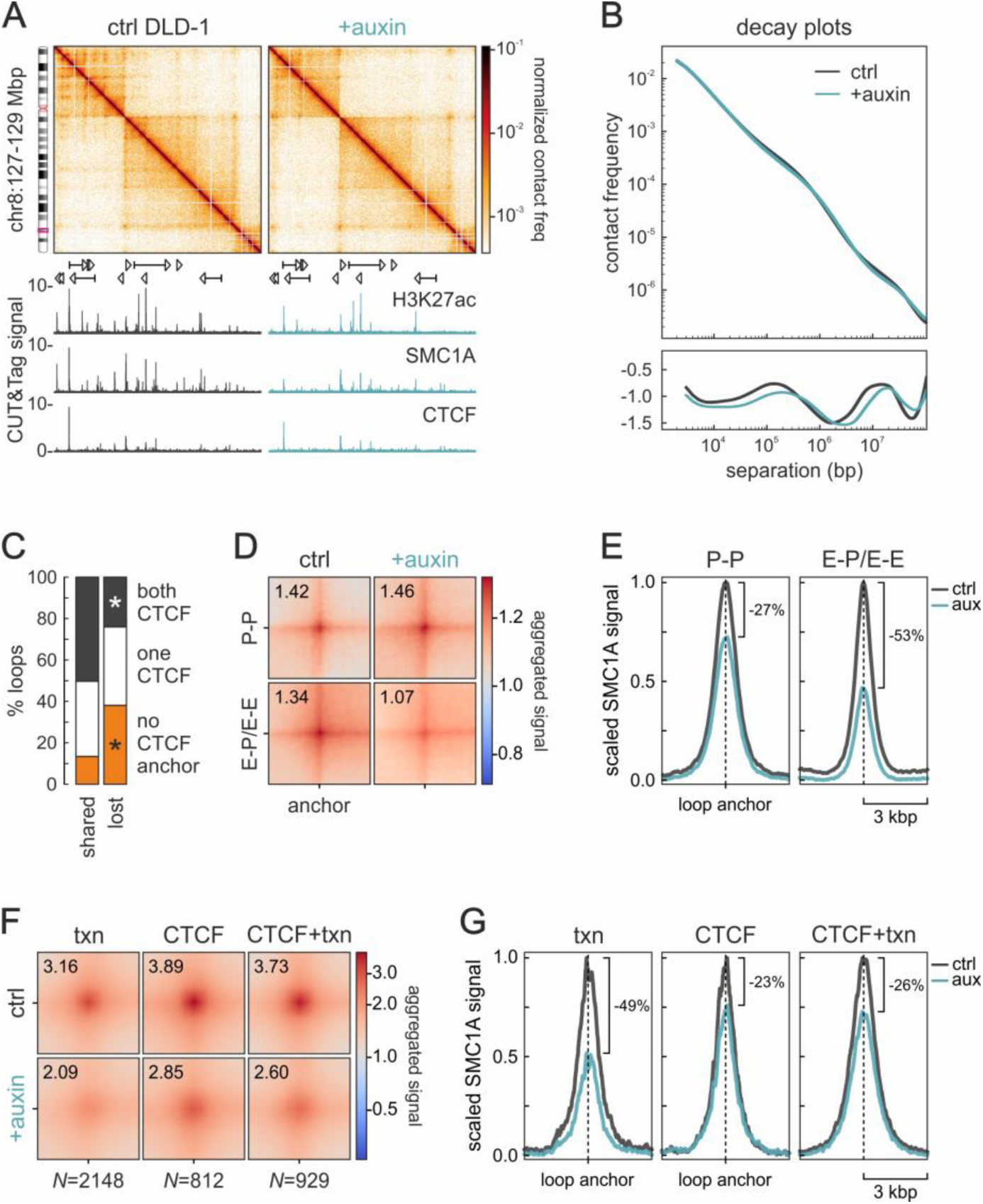
RNAPII depletion selectively affects enhancer-promoter/-enhancer loops. (**A**) Micro-C 4-kbp resolution contact maps from control (left) and auxin-treated cells (right) showing loop loss in an exemplary genomic region on chr8 aligned to H3K27ac, CTCF, and SMC1A CUT&Tag tracks. (**B**) Plots of interaction frequency decay as a function of genomic distance from control and auxin-treated cells (top) and their first derivative (bottom). (**C**) Bar plot showing per cent of loops with no (orange), one (white) or two CTCF anchors (black) that are shared between control and auxin-treated cells or lost upon RNAPII depletion. *: *P*<0.01, Fischer’s exact test. (**D**) Aggregate plots for promoter-(P-P) or enhancer-anchored loops (E-P/E-E) before (ctrl) and after RNAPII depletion (+auxin). (**E**) Line plots showing mean SMC1A signal around the loop anchors from panel D. (**F**) As in panel D, but for loops marked by transcription (txn), CTCF or by both (CTCF+txn). (**G**) As in panel E, but for the loop anchor subsets from panel F.

We also stratified loops not according to the type of elements in their anchors (i.e., enhancers *vs* promoters), but on whether their anchors were marked by CTCF, transcription or both. The majority (55%) of loops lost upon RNAPII depletion were anchored solely by RNAPII, while “CTCF-only” loops represented just 20% of the total number lost (**Figure 3F**). Notably, CTCF-bound loop anchors displayed half of the reduction in cohesin occupancy of “transcription-only” anchors (i.e., 23-26% compared to 49%; **Figure 3G**). Taken together, we discovered that RNAPII-anchored loops are more sensitive to the depletion of the polymerase, as would be predicted. However, this is selective for those loops involving enhancers in at least one anchor and not affected by the presence of CTCF.

Finally, even features of Micro-C that at first appeared unaffected upon auxin treatment respond to RNAPII depletion. For instance, loop-like signal at the edges of stripes is enhanced in the absence of RNAPII (**Figure S2E**), likely due to weakening of transcription in one of their anchors. Also, looking more carefully into loops that did not seem to change between the two conditions, we discovered that many rewired one anchor by <20 kbp (i.e., by <4 bins in a 5 kbp-resolution contact map). When we used control loop coordinates and auxin-treated Micro-C signal to plot aggregate plots, they appeared weaker. However, once we used matching auxin-treated coordinates (i.e., shifted by <4 bins in the rewired anchor) and auxin-treated Micro-C signal, we show notable strengthening of these loops too (**Figure S2F**). This suggests that depletion of RNAPII from chromatin is what also allows for such small-scale changes in loop anchor selection.

### Modeling the interplay between loop extrusion and RNAPII

A key observation in our data is the overall reduced cohesin occupancy at RNAPII loop-anchor positions (**Figures 3E**,**G** and **S2A**,**C**), which nevertheless coincides with the emergence of new prominent CTCF- anchored loops (**Figures 2A,F,G**). To test the interplay between RNAPII acting on chromatin and cohesin loop extrusion, which would be challenging to achieve experimentally, we resorted to computational modeling of 3D chromatin folding.

To this end, we considered a synthetic 460 kbp-long polymer containing two genes transcribed in the sense direction, a cluster of 8 enhancers between these two genes, plus three CTCF binding sites (two encompassing the genes and enhancers and divergently oriented to one another, and one located in the middle of the downstream gene; see **Figure 4A**). Each bead in this polymer represents 2 kbp of chromatin, and we employed two key scenarios based on established Molecular Dynamics approaches (Buckle et al., 2018; Fiorillo et al., 2020). First, a model approximating wild-type conditions (i.e., those in our untreated DLD-1 cells). In this model, RNA polymerases have specific affinity for gene promoters and enhancers, and can also transverse gene bodies at expected speeds to simulate transcription. In parallel, our model considers cohesin complexes able to bind the polymer and actively extrude loops at the experimentally-deduced speed (see **Methods** and **Table S2**). In addition, based on the documented co-association of cohesin subunits and RNAPII (Busslinger et al, 2017; Zhang et al., 2021), we introduced a weak interaction potential between cohesins and polymerases. Although cohesin is allowed to bind any position in the polymer with a probability of 0.1, it is given a 0.9 probability to bind a promoter or enhancer when found in its vicinity. In our second model, meant to approximate quantitative RNAPII depletion (i.e., like that in auxin-treated cells), RNAP molecules are removed from the simulation and cohesin is now allowed to bind any position in the polymer with equal probability.

**Figure 4.**
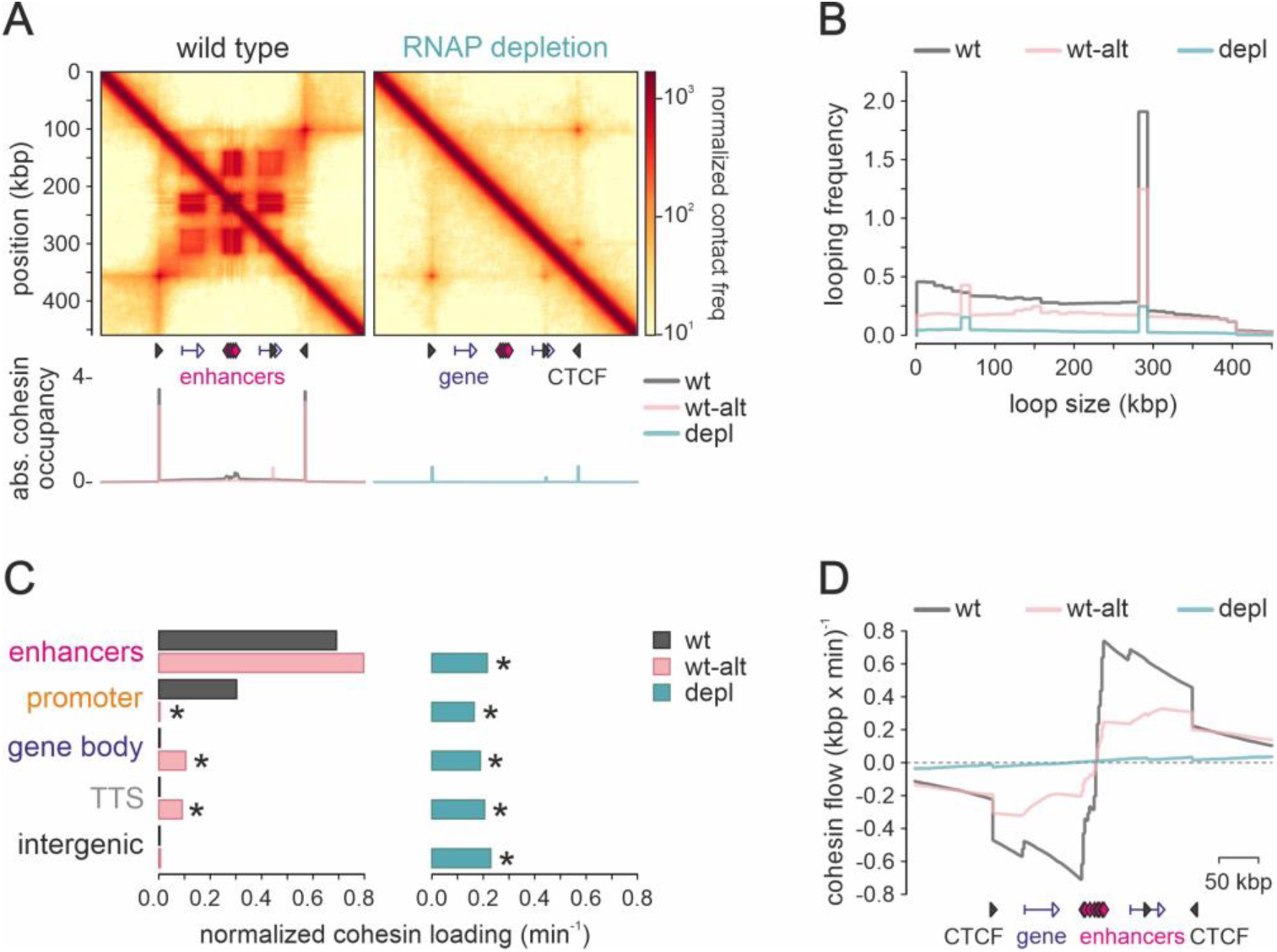
Models of 3D chromatin folding in the presence or absence of RNAPII. (**A**) Average contact maps from 1600 configurations that include (wt model) or not active polymerases (RNAP depletion) are shown aligned to plots of absolute cohesin occupancy. (**B**) Plot showing frequency of looping as a function of distance in the modeled locus under each scenario. (**C**) Bar plots showing the relative distribution of cohesin loading (per min) at different genomic elements. *: significantly different to wt; *P*<0.05, Fisher’s exact test. (**D**) Plot showing the net flow of cohesin molecules along the polymer under each scenario. Positive/negative values represent the net amount of cohesin extruding in the sense/antisense direction.

Contact maps resulting from our “wild-type” model display a ∼300-kbp CTCF-anchored loop that encompasses the two genes, and compartment-like interactions between the genes and the enhancer cluster (**Figure 4A**,**B**). Cohesin occupancy in this model shows the expected signal accumulation at the two distal, but not at the intragenic CTCF site (in line with transcribing RNAPII relocating cohesin; Heinz et al, 2019; Busslinger et al, 2017; Olan et al, 2020; **Figure 4A**). Interestingly, and despite loading favored equally at promoters and enhancers (**Figure 4C**), cohesin also accumulates at the enhancer cluster between the two genes (**Figure 4A**). Notably, this accumulation is less if we consider an “alternative wt” model where cohesin can to bind all beads with equal probability, has a weak affinity with RNAPII, but is not allowed to co-bind promoters with it. This skews loading in favor of enhancers (**Figure 4C** and as suggested by recent studies; Zhu et al., 2021; Rinaldi et al., 2022; Rinzema et al., 2022). In this “wt-alt” model (**Table S2**), we also see increased cohesin presence and loop formation at the intragenic CTCF (**Figure 4B**; contact map not shown), which does not agree with our Micro-C data. Nevertheless, in both models transcription start sites act as potent boundaries to cohesin-driven extrusion (**Figure 4A,D**; as also eluded to by Preprint: Valton et al., 2021).

Implementation of the “RNAP depletion” model leads to the elimination of all contacts between genes and promoters in parallel with the emergence of a new CTCF loop anchored at the intragenic CTCF site (**Figure 4A**). This is in agreement with our experimental findings of intragenic CTCF sites being disproportionately engaged in looping following RNAPII degradation (**Figure 2F,G**), as well as with the accumulation of cohesin at the three CTCF-bound sites in our model (**Figure 4A,B**). Thus, we can deduce that it is the competition between transcribing RNAPs and cohesin loaded preferentially, but not exclusively, at promoters and enhancers that gives rise to the contact patterns mapped via Micro-C. However, our models cannot yet explain the selective loss of enhancer-promoter interactions following RNAPII depletion, as both elements are modeled using similar general attributes.

## DISCUSSION

In previous work, we established the necessity of RNAPII for refolding interphase chromatin during the mitosis-to-G1 transition of a human cell line. We did not only show domain and compartment erosion in the absence of RNAPII, but also a dependency for cohesin loading onto chromatin (Zhang et al., 2021). However, in that same study, we could not identify 3D architecture changes of comparable magnitude using Hi-C from non-synchronized RNAPII-depleted G1 cells. This remained perplexing until we obtained the Micro-C data analyzed here. Our highly-resolved interaction maps unveil a transcription-based architecture of interphase chromatin that is markedly perturbed by RNAPII depletion. We documented both loss and gain of specific loop-like spatial interactions that beg the following questions.

First, how do longer and more pronounced CTCF loops arise in the absence of RNAPII? As RNAPs are thought to represent physical barriers for cohesin and transcriptional elongation to antagonize loop extrusion (Rowley et al., 2019), depletion of RNAPII would allow for a more efficient extrusion of loops anchored at CTCF-bound sites. This can be considered the converse of recent observations whereby transcribing RNAPII complexes reposition cohesin complexes (Busslinger et al., 2017; Heinz et al., 2018; Olan et al., 2020) or give rise to new spatial interactions (Rosencrance et al., 2020). Hence, the *de novo* engagement of CTCF sites located inside of previously-active promoters and gene bodies predominates the anchors of loops forming upon RNAPII dpeletion (as also corroborated by our simulations). Of note is the subset of loops emerging in the absence of RNAPII via interactions between Polycomb-bound regions. Although these H3K27me3-marked regions are considered transcriptionally inert, they often bind “poised” RNAPII (Ferrai et al., 2017), removal of which might contribute directly or indirectly to the observed effects. Directly by the competitive interplay of RNAPII with Polycomb complexes, and indirectly by RNAPs affecting cohesin loading (as cohesin depletion was shown to enhance interactions between Polycomb-bound regions; Rhodes et al., 2020). A somewhat similar effect is now described for promoter-proximal paused RNAPs maintaining local 3D chromatin architecture in differentiated erythrocytes (Preprint: Penagos-Puig et al., 2022).

Second, why do promoter- and enhancer-anchored interactions respond differently to RNAPII depletion? Here, we recorded two unforeseen events. H3K27ac levels drop genome-wide upon RNAPII depletion, but are significantly more reduced at enhancers compared to promoters. At the same time, enhancer-anchored loops are selectively weakened upon RNAPII depletion, but promoter-promoter ones remain unaffected. This is independent on whether these interactions have or not CTCF at their anchors, and reflected on cohesin occupancy decreasing in a manner similar to H3K27ac. This is striking, given that promoters and enhancers are thought to be subtle variants of a single class of *cis*-elements (Andresson et al., 2015). Nevertheless, our data suggest that spatial communication among promoters relies on a different set of factors that the communication involving enhancers only or enhancers and their target promoters. Which these factors and their attributes are is the next challenge in this work.

Third, how is the presence of cohesin on chromatin affected in the absence of RNAPII? In the M-to-G1 transition, we could correlate reduced cohesin loading to the depletion of RNAPII from chromatin despite the fact that chromatin accessibility was not reduced (and, thus, could not be solely responsible for any reduction in cohesin loading; Zhang et al., 2021). Now, we document a comparable reduction in interphase nuclei upon RNAPII depletion, which is most apparent at enhancers-promoter loops. This is in line with the binding of cohesin loaders at gene promoters (Busslinger et al., 2017, although the specificity of some of this data is now debated; Preprint: Banigan et al., 2022); with the fact that the loader NIPBL and unloader WAPL co-purify with RNAPII complexes (Zhang et al., 2021); as well as with recent work pointing to enhancers as cohesin-loading sites for the formation of 3D interactions (Liu et al., 2021a; Zhu et al., 2021; Rinaldi et al., 2022; Rinzema et al., 2022). Moreover, we should reconsider studies where pioneer transcription factors (like OCT4 and SOX2; Liu et al., 2021a) and chromatin remodelers (like SNF2h; Hakimi et al., 2002) affected the loading/unloading cycles of cohesin onto chromatin. Still, distinguishing between cohesin loading or stalling at a given position *in vivo* remains challenging, due to the highly processive nature of extruding complexes. However, our simulations show that, by introducing a weak interaction between RNAPII and cohesin, the latter is predominantly directed to active promoters and enhancers. Disfavoring cohesin loading at promoters (by simulating competition by RNAPII), generates contacts rarely seen in Micro-C data.

Cohesin, its loader NIPBL, and the Mediator complex were proposed to co-associate in order to physically and functionally connect active enhancers and promoters (Kagey et al., 2010). Hi-C studies that followed this early work found Mediator and RNAPII to be dispensable for 3D chromatin folding (Jiang et al., 2020; El Khattabi et al., 2019). This is now challenged by MCC data showing loss of enhancer-promoter contacts following acute depletion of Mediator (Preprint: Ramasamy et al., 2022) and by decreased cohesin binding to RNAPII-transcribed genes following depletion of the yeast Med14 subunit (Mattingly et al., 2022). These new observations, together with the Micro-C analysis we present here, put the still-debated role of RNAPII in 3D chromatin folding in a different light. Especially given that RNAPII depletion does not cause a commensurate loss of Mediator from chromatin (**Figure S1A**).

In summary, we now present definitive evidence for the involvement of RNAPII in sustaining enhancer-promoter interactions, as well as in the direct competition towards the formation of CTCF loops. The former appears to require the presence of RNAPII on chromatin, whereas the latter likely also implicates ongoing transcription. Nonetheless, there remain aspects of this RNAPII-based chromatin architecture to be elucidated, like the differential dependency of gene promoter-compared to enhancer-anchored interactions or the mechanistic details of its influence on cohesin loading.

## ACKNOWLEDGEMENTS

We thank Marieke Oudelaar, Kerstin Wendt, and all members of the Papantonis lab for discussions, and the Maeshima lab (NIG, Japan) for the DLD-1 mAID-RPB1 cells.

## Author contributions

S.Z. performed bioinformatics analyses. N.Ü. performed experiments. M.B. performed computational modeling. A.P. conceived and supervised the study, and compiled the manuscript with input from all coauthors.

## Data availability

NGS data generated in this study are available via the NCBI Gene Expression Omnibus repository under the accession number GSE178593.

## METHODS

### Cell synchronization and sorting

mAID-POLR2A(RPB1)-mClover DLD-1 cells (Nagashima et al., 2019) were grown in RPMI-1640 medium supplemented with 10% FBS under 5% CO2. Inducible depletion of RPB1 initiated via treatment with doxycycline for 24 h to induce *TIR1* expression, before addition of 500 µM indole-3-acetic acid solution (“auxin”, Sigma-Aldrich) for 14 h to induce RPB1 degradation. Cells treated with auxin were harvested, resuspended in 1 µg/ml propidium iodide, and sorted to isolate G1 cells on a FACS Canto II flow cytometer (Becton Dickinson).

### Micro-C and data analysis

Micro-C was performed using the Micro-C v1.0 kit in collaboration with Dovetail Genomics as per manufacturer’s instructions. Micro-C libraries (at least 3 per each biological replicate) that passed QC criteria were pooled and paired-end sequenced on a NovaSeq6000 platform (Illumina) to >600 million read pairs per replicate (**Table S1**). Micro-C contact matrices were produced using Dovetail Genomics pipeline (https://micro-c.readthedocs.io/en/latest/fastq_to_bam.html). In brief, read pairs were mapped to human reference genome hg38 using BWA, after which low mapping quality (<40) reads and PCR duplicates were filtered out. Next, ICE-balanced .*cool* files and KR-balanced .*hic* files were generated and visualized via HiGlass or Juicebox. Decay plots were generated *cooltools* (https://cooltools.readthedocs.io/en/latest/notebooks/contacts_vs_distance.html). Subcompartment analysis was performed using CALDER (Liu et al., 2021b) considered 50 kbp-resolution Micro-C data. For loop calling, we used a multi-tool (HiCCUPS, *cooltools*, and *mustache*) and multi-resolution (5- and 10-kbp) approach as previously described (Hsieh et al., 2020; Krietenstein et al., 2020). Loop lists derived from each tool were merged using *pgltools* (Greenwald et al., 2017) as follows: dots from both 10- and 5-kbp resolution are retained if they are supported by >10 read counts, and kept at native resolution. To further annotate loops as CTCF- or transcription-anchored, using CTCF, H3K27ac CUT&Tag peaks (from this work), as well as RNAPII peaks and nascent RNA-seq signal (RPKM >10; from Zhang et al., 2021). All intersections were performed using *pgltools intersect1D* without any distance tolerance for CTCF anchors, and with a 10-kbp tolerance for enhancers and promoter anchors (annotated TSS±2 kbp) identified using *chipseeker* (Yu et al., 2015). Note that promoters of all gene isoforms were considered, and “super-enhancers” called using the ROSE algorithm (Whyte et al., 2013). Finally, aggregate plots for loops and boundaries were generated using *coolpup*.*py* (Flyamer et al., 2020). Code used in this study is available at: https://github.com/shuzhangcourage/HiC-data-analysis.

### Cleavage Under Targets and tagmentation (CUT&Tag)

Following lifting from plates using accutase and FACS sorting, 0.5 million G1-phase DLD-1 cells were processed according to manufacturer’s instructions (Active Motif). Samples were paired-end sequenced to obtain at least 10^7^ reads. Reads were processed according to the standard CUT&Tag pipeline (https://yezhengstat.github.io/CUTTag_tutorial/). Briefly, paired-end reads were trimmed for adapter removal and mapped to human (hg38) and *E. coli* reference genomes (ASM584v2) using Bowtie 2 (Langmead & Salzberg, 2012). *E. coli* mapped reads were quantified and used for calibrating human-mapped reads. Peak calling was performed using a multi-FDR-tryout method (FDR <0.01 to <0.1) and IgG controls for thresholding. Acceptable FDRs could vary between different datasets, but were always kept same for control and auxin-treated samples. Thus, for CTCF, an FDR <0.1 was selected and, for additional stringency, we only considered a CUT&Tag peak as CTCF-bound if it encompassed a canonical CTCF motif (assessed via fimo; Charles et al., 2011). For H3K27ac and H3K27me3, peaks were selected on the basis of FDR <0.025 and <0.01, respectively, while for SMC1A, an FDR <0.1 was used. Heatmaps were generated using Deeptools (Ramirez et al., 2014).

### Chromatin fractionation and western blotting

For assessing protein abundance in different subcellular fractions, a protocol previously described was used (Watrin et al., 2006). Protein concentration in each fraction extract was determined using the Pierce BCA Protein Assay Kit (Thermo Fisher Scientific). Following separation on precast SDS-PAGE gels (BioRad), proteins were detected using antibodies against the RNAPII-CTD (Abcam, ab817; 1:500), phospho-Ser5 RPB1 (Active Motif #61085; 1:2000), MED24 (Affinity Biosciences, AF0346; 1:1000), Lamin B1 (Abcam, ab16048; 1:10000), and HSC70 (Santa Cruz, sc-7298; 1:2000), and visualized using the Pierce SuperSignal West Pico ECL kit (Thermo Fisher).

### Simulations of chromatin folding

We performed Molecular Dynamics simulations via the multi-purpose EspressoMD package (Reynwar et al., 2007). In our simulations, individual proteins are represented by ‘‘beads’’ interacting via phenomenological force fields and move according Langevin equation, and the chromatin fiber is represented as a chain of beads connected by bonds. The position of every bead in the system, either a protein or chromatin bead, evolves according to the Langevin differential equation that encodes Newton’s laws in the case of thermal bath with the friction *γ* due to an implied solvent in presence of forces between beads encoded by energy potential functions *U* (Chiariello et al., 2016; Buckle et al., 2018). Langevin equations for all beads are simultaneously solved in EspressoMD using a standard Velocity-Verlet numerical algorithm. The potential connecting *i* and *i+1* beads of the fiber is a finitely extensible non-linear elastic (FENE) spring that adds up to a steric repulsion potential between non-adjacent sites of the polymer, the Weeks-Chandler-Andersen (WCA) potential:

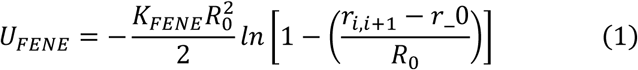

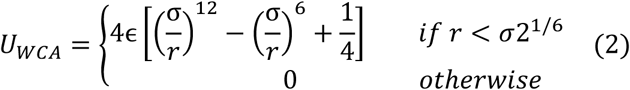

where *r*_*i,i*+1_ is the distance between consecutive beads, and σ is where the interaction from repulsive becomes attractive and can be interpreted as the diameter of the particles. This value is a natural length scale of the system. In FENE we fix parameters to have an equilibrium distance of 1.6 σ with maximum extension of 0.8 σ, and a bond energy of *K*_*FENE*_ = 30 k_B_T. Since our fiber is resolved at 2 kbp, chromatin rigidity cannot be neglected (i.e. that we are below the estimated persistence length). Bending rigidity of the polymer is introduced via the Kratky-Porod potential for every three adjacent chromatin beads where θ is the angle between three consecutive beads as given by:

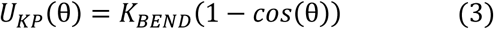

and K_BEND_ is the bending energy. The persistence length in units of σ is given by *L*_*p*_ *= K*_*BEND*_*/k*_*B*_*T*.

In order to model more complex aspects of transcription and loop extrusion, their interplay and impact on 3D chromatin organization, we encoded in the model: (i) full 3D loop extrusion by interplay of cohesin dimers and CTCF, and (ii) transcription by RNAPII particles. To simulate association of cohesin and RNAPII with the chromatin fiber, we employed a harmonic potential mimicking formation of a stable bond between two particles that fluctuate around an equilibrium distance d_0_:

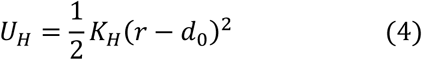

Whenever in the following description the formation of a bond is referred to, we mean the introduction of the aforementioned harmonic bond between the particles involved. To regulate the lifetime of the such a harmonic interaction, we introduced mechanisms of bond formation and removal according to cutoff distance *c*_*d*_ below which a bond is formed, with a certain probability rate of detachment in units of time τ_b_=2 τ, τ the fundamental MD unit of time (see below). These are then set to approximate the experimentally-observed range of RNAPII transcription and cohesin loop extrusion speeds and chromatin residence time. Introduction of the above mechanics is added on top of the SBS model we previously employed (Barbieri et al., 2017). The SBS model encodes the association tendency of RNAPII with promoters by means of the shifted, truncated Lennard-Jones (LJ) potential that allows spontaneous co-localization of beads around a distance σ with lifetime and stability properties depending on the depth of the energy well ϵ:

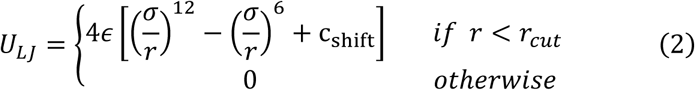

where *r*_*cut*_ is a cutoff distance, *r* is the separation of any two beads, and *r*_*cut*_ = 2.5σ for all LJ potentials in the simulations. This is a standard widely used in the field to simulate phenomenological course-grained affinities (Chiariello et al., 2016; Buckle et al., 2018).

For RNAPII interactions and transcription, the polymerase is represented as a bead with LJ interaction with specific beads of the chromatin fiber representing promoters, enhancers or gene body with energy ϵ = 3 k_B_T. Such relatively mild affinity helps to identify promoters as the correct sites where transcription initiation will take place (i.e., RNAPII forming stable bonds with promoter beads) before the elongation process along the gene body starts. LJ interactions were also introduced among RNAPII beads (ϵ=3.5k_B_T) to simulate their tendency to form condensates acting as transcription hubs, as well as between RNAPII and cohesin (ϵ=3k_B_T) to simulate its preferential loading at promoter/enhancer beads. RNAPII transcription dynamics are simulated as a four-step process: attachment to a promoter in an exclusive manner, elongation starts, elongation proceeds through the gene body, detachment at TES. A bond forms if the beads are less distant in space than a certain cutoff (2.8σ). To simulate the tendency of RNAPII to reel in gene body beads, another secondary bond is formed with the next bead on the chromatin fiber in the direction of transcription (*i+1* bead, where *i* is the promoter coordinate on the fiber and if transcription occurs in the sense direction). In the next step, for RNAPII to move on the *i+1* site, a new bond needs to form between the two, the old bond needs to dissolve, and a new secondary bond to form with *i*+2 bead. This happens at a given rate (0.4 τ_b_^-1^) and only if the beads are found within less than a cutoff distance (c_d_=1.05σ). All following steps along the gene body (third step) do not require an acceptance rate, but to only satisfy the cutoff distance. Every time a new bond forms, the previous one dissolves. Upon reaching the TES, RNAPII stops and at a given rate (0.2 τ_b_^-1^) can become unbound. When transitioning from the unbound to the promoter-bound state, RNAPII loses its LJ interaction with promoters and gene body, since this is substituted by the bond itself. However, RNAPII retains the LJ interaction with enhancers, since it is not allowed to form bonds with enhancer beads or with other unbound RNAPIIs. Such retained interactions are increased in magnitude (from 4 to 6 k_B_T) to favor associations with actively transcribed segments. This preserves the stability of hubs/ condensates during the process of transcription.

For CTCF binding, a single bead interacting via LJ interactions (ϵ=3k_B_T) with specific beads of the chromatin fiber representing oriented cognate binding motifs. Once a bond forms, at a certain rate (0.8 τ _b_ ^-1^) it is exclusive (i.e., other CTCF cannot bind that same site). This binding dissolves at a given rate (10^−4^ τ _b_^-1^) and CTCF is again free to diffuse and search for other binding sites. On the other hand, cohesin is represented as a pair of beads connected by a bond (r_0_=1.6σ and K=8 k_B_T). Extrusion has three steps: attachment, active extrusion, and detachment. For attachment, each cohesin monomer forms a bond with the chromatin fiber. Bonds form when a cohesin monomer and a chromatin site come within a given cutoff distance (1.6σ) and at certain rate (0.1 τ_b_^-1^). Only the case where both monomers simultaneously form bonds on adjacent chromatin beads is considered a successful attachment and the dimer is retained for the next step, otherwise bonds dissolve. To give monomers and the next bead (*i±1* bead on polymer, where *i* is the attachment site) the tendency to come closer and start extrusion, same way as RNAPII transcription above, a new secondary bond forms at the time of attachment with the i±1 bead in the direction of extrusion. Hence, extrusion consists of creating new bonds with the *i±1* and *i±2* beads, while dissolving the old bonds. Also in this case, new bonds only form if the distance is below a cutoff value (1.1σ). Such value produces ranges of cohesin extruding speed of 12-16 kbp/min, which is within the range of experimentally observed values (Rao et al., 2017). This allows cohesin to be affected by the surroundings during extrusion (e.g., by the presence of RNAPII close by or bound to adjacent sites). Finally, cohesin detachment can occur independently at any step at a given rate (10^−4^τ_b_^-1^) to fit its known chromatin residence time of 20 min. Then, CTCF “loop anchors” are modeled such that it is impossible for cohesin to form new bonds with the next *i±1* site if the latter is already bound by CTCF, provided it has the binding motif in convergent orientation. This renders extrusion dependent on CTCF dynamics. Last, cohesin also has LJ potential for interacting with RNAPII both in the bound (ϵ=6 k_B_T) and unbound state (ϵ=3 k_B_T) with a preference for bound RNAPII to its suggested role in cohesin loading on the chromatin fiber (Zhang et al., 2021). RNAPII and LE dynamics are performed using a python script that drives the EspressoMD library. The polymer initializes as a random walk and its dynamics first evolves in the absence of extrusion and transcription to generate an equilibrium coil conformation. In the following step, both extrusion and transcription are switched on, and its dynamics evolve until a new steady-state conformation is obtained. Across all simulations, we used standard values for the friction coefficient (*γ*=0.5) and the time step (t=0.01), and we let the system evolve for up to 10^8^ steps. As in previous studies, to connect our *in silico* space-time units with real distances and times of the biological process, we assumed that the concentration of DNA in the 3D simulation space is the same as that in a human nucleus. If we use a total DNA amount of 6 Gbp and a nucleus radius of 5 μm, we obtain the rough estimation of σ=64 nm. For time units, we consider the standard MD relation τ=η(6 π σ^3^/ϵ). Assuming a viscosity ∼0.5P, the fundamental time unit is τ=0.06 sec. By running simulations starting from independent configurations and by sampling periodically the system we obtain an ensemble of configurations up to 5×10^3^ for the measurement of the quantities shown. Concentrations of CTCF, cohesin, and RNAPII are taken from physiological values and range from 75 to 300 nmol/l. The energy scale of the system is given by the Boltzmann factor kB multiplied by the temperature of the system T=310 K.

### Statistical analyses

*P*-values derived from the Fisher’s exact test were calculated using GraphPad (http://graphpad.com/), those from the Wilcoxon-Mann-Whitney test using R. Unless otherwise stated, *P*-values <0.01 were deemed significant.

## SUPPLEMENTARY MATERIAL

**Figure S1.**
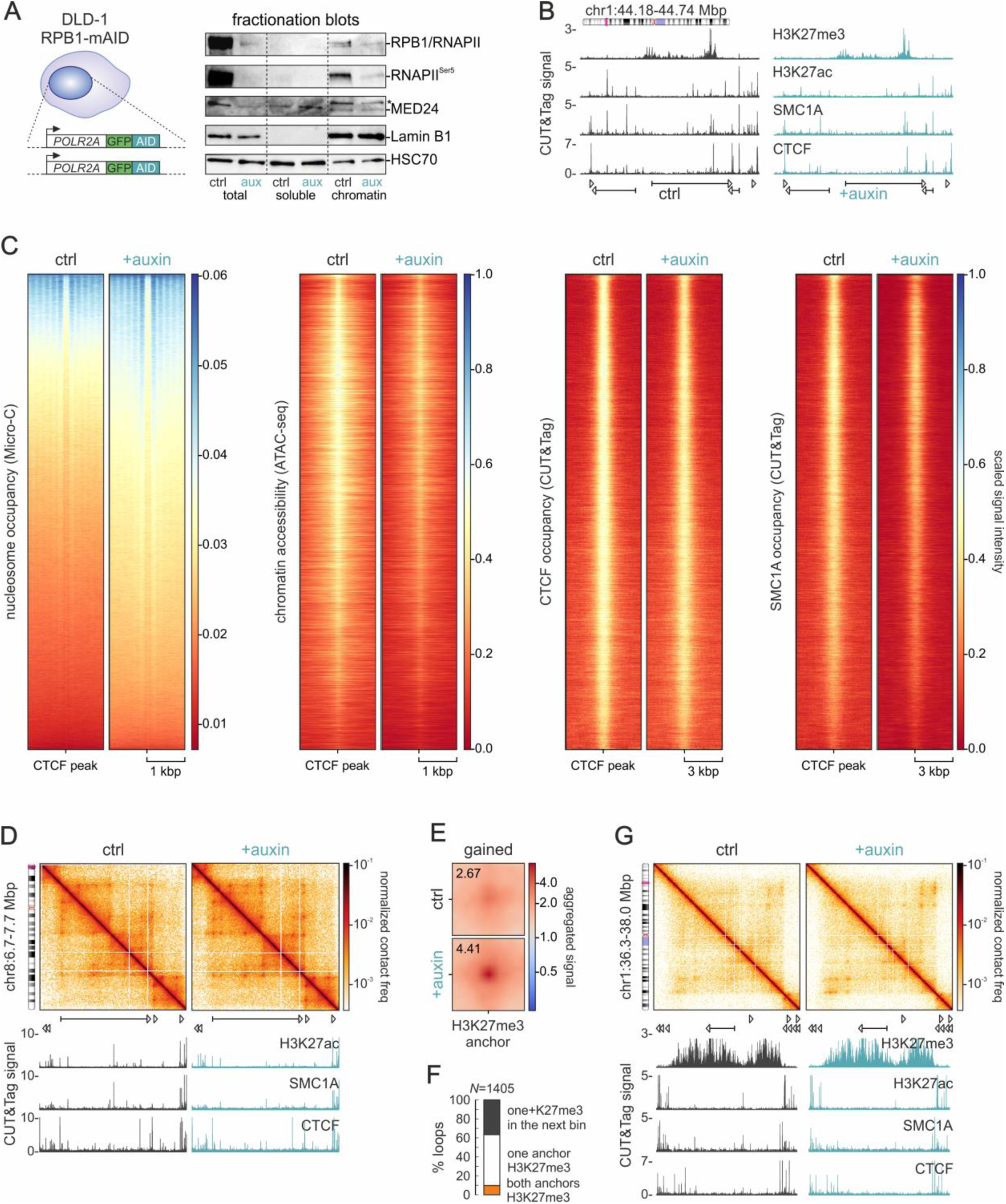
Effects of RNAPII depletion on eu- and heterochromatin 3D organization. (**A**) Left: Schematic of the bi-allelic tagging strategy in the endogenous *POLR2A* locus. Right: Fractionation blots showing the levels of RPB1 and Ser5-phosphorylated RNAPII, Mediator subunit 24, and Lamin B1 from DLD1-mAID-RBP1 cells treated or not with auxin to deplete RNAPII. HSC70 levels provide a control. (**B**) Representative tracks of CUT&Tag signal for H3K27me3, H3K27ac, SMC1A, and CTCF from control (*black*) and auxin-treated DLD1-mAID-RBP1 cells (*green*) along 0.55 Mbp of chr1. (**C**) Heatmaps showing nucleosome occupancy deduced from Micro-C data, chromatin accessibility deduced from ATAC-seq, and CTCF and SMC1A occupancy around CTCF loop anchors before (ctrl) and after RNAPII degradation (+auxin). (**D**) Micro-C contact maps from control (left) and auxin-treated cells (right) in an exemplary genomic region on chr8 at 2-kbp resolution aligned to H3K27ac, CTCF, and SMC1A CUT&Tag tracks. (**E**) Aggregate plots of the 890 H3K27me3-anchored loops emerging in auxin-treated cells. (**F**) Bar plot showing per cent of gained loops with one (white) or two H3K27me3 anchors (orange) or with H3K27me3 in the next genomic bin (i.e., within <10 kbp from the anchor). (**F**) As in panel D, but in a region of chr1 where H3K27me3-anchored loops form upon RNAPII depletion.

**Figure S2.**
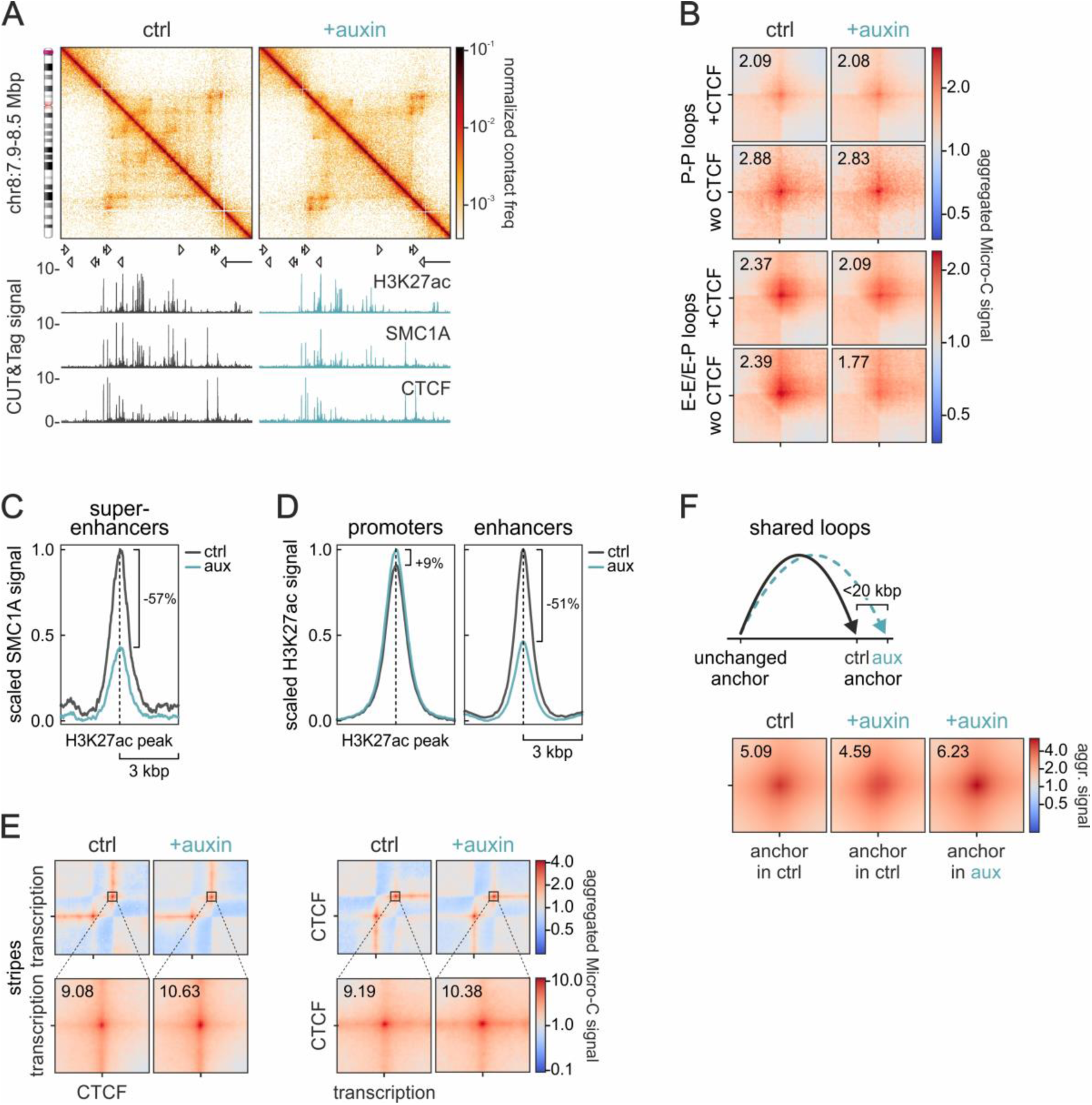
Properties of loops gained upon RNAPII depletion. (**A**) Micro-C contact maps from control (left) and auxin-treated cells (right) in an exemplary genomic region on chr8 at 4-kbp resolution aligned to H3K27ac, CTCF, and SMC1A CUT&Tag tracks. (**B**) Aggregate plots of promoter-(P-P) or enhancer-anchored loops (E-E/-P) in control and auxin-treated cells that involve (+CTCF) or not CTCF (wo CTCF) in at least one anchor. (**C**) Line plot showing mean SMC1A signal from control and auxin-treated cells in the 6 kbp around H3K27ac peaks in 590 super-enhancers. (**D**) As in panel C, but for H3K27ac signal around active promoters or enhancers. (**E**) Average plots showing mean signal of stripes with one CTCF and one transcriptional anchor before (ctrl) and after RNAPII depletion (+auxin). *Zoom-in*: Aggregate plots for loops at the end of stripes. (**F**) As in panel B, but for shared loops that rewire one anchor by <20 kbp.

**Table S1.** Micro-C statistics. Mapping, deduplication, and contact statistics from two merged replicates.

|  | Control DLD-1 | % | +Auxin DLD-1 | % |
| --- | --- | --- | --- | --- |
| Total read pairs | 1,648,350,580 | 100.0 | 1,524,930,442 | 100.0 |
| Mapped read pairs | 1,355,832,740 | 82.25 | 1,249,138,654 | 81.91 |
| Duplicate pairs | 448,059,906 | 27.18 | 402,268,327 | 26.38 |
| Valid *trans* read pairs | 103,889,160 | 11.44 | 102,355,674 | 12.09 |
| Valid *cis* read pairs | 803,883,674 | 88.56 | 744,514,653 | 87.91 |
| Long-range contacts | 559,791,780 | 61.67 | 507,103,457 | 59.88 |
| Long-/short-contact ratio | 2.29x | N/A | 2.14x | N/A |

**Table S2.**
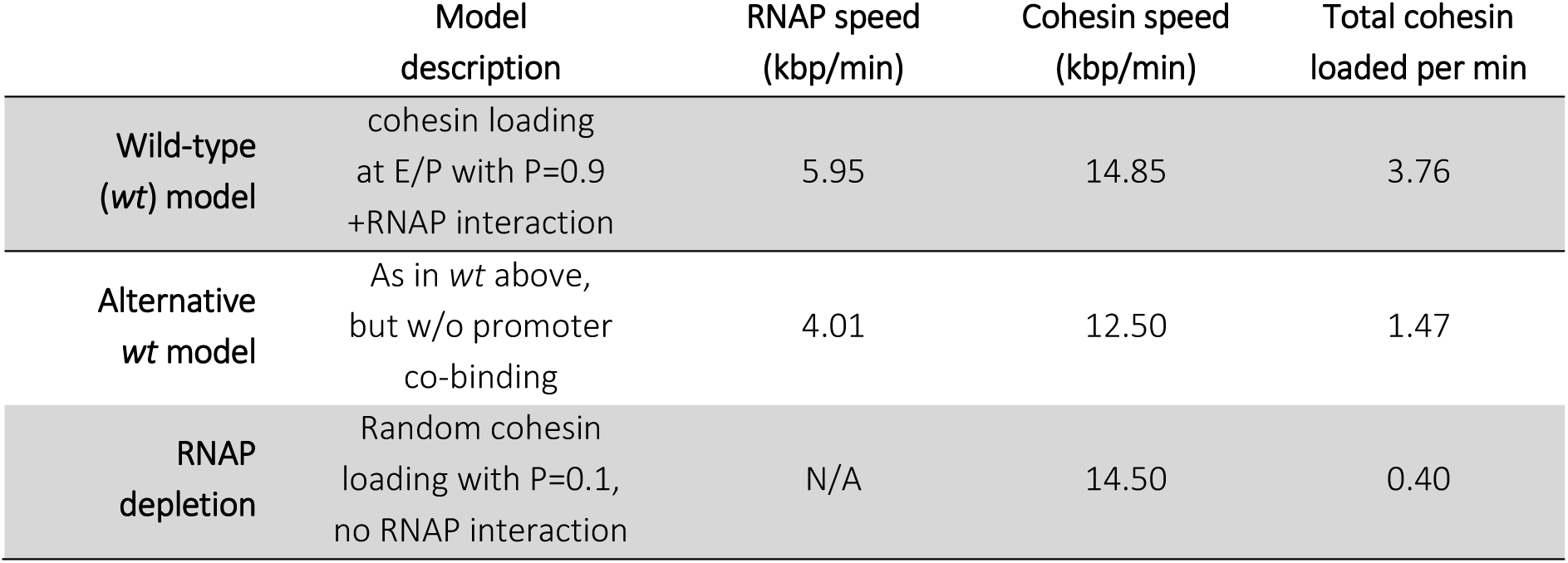
Simulation parameters. Translocation speeds and loading in the different MD scenarios.

